# Centriole distal-end proteins CP110 and Cep97 influence centriole cartwheel growth at the proximal-end

**DOI:** 10.1101/2021.07.08.451650

**Authors:** Mustafa G. Aydogan, Laura E. Hankins, Thomas L. Steinacker, Mohammad Mofatteh, Saroj Saurya, Alan Wainman, Siu-Shing Wong, Xin Lu, Felix Y. Zhou, Jordan W. Raff

## Abstract

Centrioles are composed of a central cartwheel tethered to nine-fold symmetric microtubule (MT) blades. The centriole cartwheel and MTs are thought to grow from opposite ends of these organelles, so it is unclear how they coordinate their assembly. We previously showed that an oscillation of Polo-like kinase 4 (Plk4) helps to initiate and time the growth of the cartwheel at the proximal end. Here, we show that CP110 and Cep97 form a complex close to the distal-end of the centriole MTs whose levels rise and fall as the new centriole MTs grow, entrained by the core Cdk/Cyclin oscillator that drives the nuclear divisions in these embryos. These CP110/Cep97 dynamics, however, do not appear to time the period of centriole MT growth directly. Instead, we find that changing the levels of CP110/Cep97 alters the Plk4 oscillation and the growth of the cartwheel at the proximal end. These findings reveal an unexpected crosstalk between factors normally concentrated at opposite ends of the growing centrioles, which may help to coordinate centriole growth.

## Introduction

The cytoplasm is a compact environment, filled with many different types of organelles that often have structurally complex conformations. How these organelles coordinate their assembly in a precise and timely manner is a fundamental question in cell biology (Liu et al., 2018; Mukherji and O’Shea, 2014). Centrioles are cytoskeletal organelles whose linear structure makes them an excellent model with which to study the principles of organelle biogenesis. In most dividing cells, centrioles duplicate in S-phase when a daughter centriole assembles from the side of an existing mother centriole (Arquint and Nigg, 2016; Banterle and Gönczy, 2017; Fırat-Karalar and Stearns, 2014). Newly formed centrioles are composed of an inner cartwheel and surrounding microtubule (MT) blades – two structures that are thought to be assembled at opposite ends of the growing daughter centriole (Breslow and Holland, 2019; Gemble and Basto, 2018). In many species, the cartwheel and MT blades grow to approximately the same size (Winey and O’Toole, 2014), indicating that the growth of the centriole cartwheel and centriole MTs are coordinated to ensure that these structures grow to consistent absolute and relative sizes. There is also a second phase of centriole growth that is thought to occur largely in G2, and in some species or cell-types this can be substantial, so that the centriole MT blades extend well beyond the cartwheel; this second phase of growth may be less tightly regulated (Kong et al., 2020).

We recently showed in early *Drosophila* embryos that cartwheel assembly is homeostatic: when cartwheels grow slowly, they grow for a longer period, and *vice versa* to maintain a constant size (Aydogan et al., 2018). Plk4 helps to enforce this homeostatic behaviour by forming an oscillating system at the base of the growing cartwheel that can be entrained by the core Cdk/Cyclin cell-cycle oscillator (CCO) to help initiate and time cartwheel formation in S-phase (Aydogan et al., 2020). In contrast, we know relatively little about how the growth of the centriole MTs is regulated (Sharma et al., 2021). Proteins such as Sas-4/CPAP (Schmidt et al., 2009; Sharma et al., 2016; Zheng et al., 2016), Cep135/Bld10 (Dahl et al., 2015; Lin et al., 2013) and Ana1/Cep295 (Alvarez-Rodrigo et al., 2021; Chang et al., 2016; Saurya et al., 2016) are thought to interact with the centriole MTs to promote centriole MT growth, while proteins such as CP110 and Cep97 are concentrated at the distal-end of the centrioles where they appear to suppress centriole MT growth (Delgehyr et al., 2012; Franz et al., 2013; Kohlmaier et al., 2009; Schmidt et al., 2009; Spektor et al., 2007). It is unclear, however, how these proteins are regulated to ensure that the centriole MTs grow to the correct size, nor how this growth is coordinated with the growth of the centriole cartwheel.

Here we show that in early fly embryos the levels of CP110 and Cep97 rise and fall at the distal end of the growing daughter centriole over the course of a nuclear division cycle. The gradual accumulation of CP110/Cep97 at the tip of the growing daughter centriole, however, does not appear to set the period of daughter centriole MT growth. Instead, we find that CP110 and Cep97 levels in the embryo influence the growth of the centriole cartwheel, apparently by tuning the Plk4 oscillation at the base of the growing daughter. These findings reveal that crosstalk between factors normally localized to opposite ends of the growing centrioles could help to coordinate centriole cartwheel and MT growth.

## Results

### CP110 and Cep97 form a complex close to the (+) end of centriole MTs

To examine the centriolar localization of CP110 and Cep97 in *Drosophila*, we generated transgenic fly lines expressing either CP110-GFP or GFP-Cep97 under the control of the *Ubiquitin* (*Ubq*) promoter (uCP110-GFP and uGFP- Cep97) that also expressed Asl-mCherry (as a marker for the mother centriole) (Novak et al., 2014). The GFP-fusions are moderately overexpressed (by ∼2-5X; estimated by blots of serial dilutions of the extracts) compared to the endogenous protein (Franz et al., 2013) (Fig. S1A; see also Fig.4, A and B). In fixed wing-disc cells, 3D-Structured Illumination super-resolution Microscopy (3D-SIM) confirmed previous observations that CP110/Cep97 localize to the distal-end of the mother and daughter centrioles (Dobbelaere et al., 2020; Franz et al., 2013) (Fig. 1A). Moreover, the resolution of this system in fixed cells was sufficient to reveal that both proteins assembled into a ring at the distal end of the mother centriole that is of the correct size to be positioned at, or very close to, the plus-ends of the centriole MTs (Fig. 1) – consistent with previous reports (LeGuennec et al., 2020; Yang et al., 2018).

**Figure 1.**
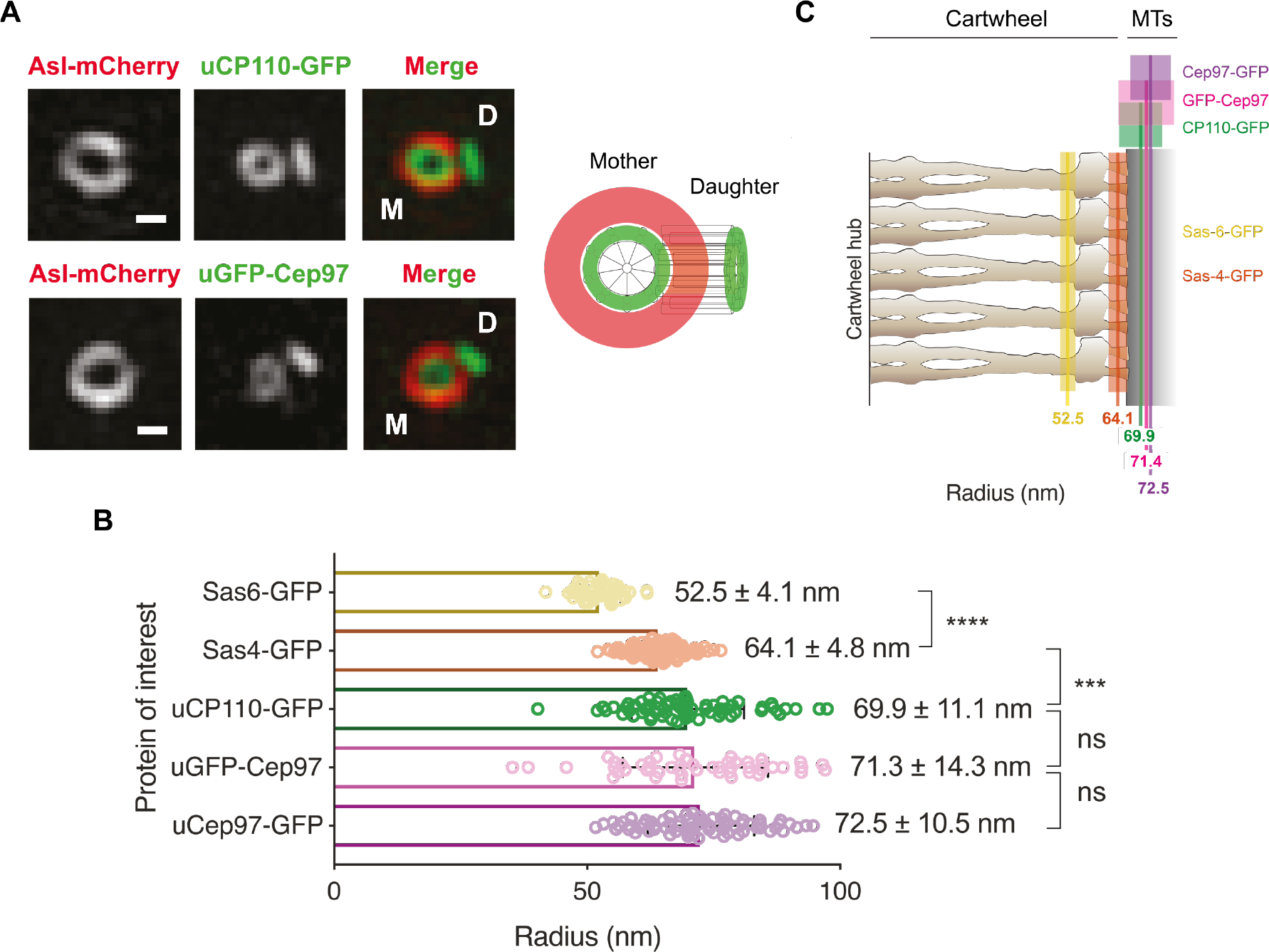
3D-SIM analyses reveal that CP110 and Cep97 co-localise near the distal-end of the centriole microtubules. **(A)** 3D-SIM micrographs show the distribution of the mother centriole marker Asl-mCherry and either uCP110-GFP or uGFP-Cep97 at mother/daughter centriole pairs in *Drosophila* wing disc cells (Scale bars=0.2 μm). uCP110- GFP and uGFP-Cep97 are located at the distal ends of the mother and daughter centrioles (with the mother centriole viewed “end-on”, as depicted in the schematic). **(B)** The horizontal bar chart quantifies the mean radii of the indicated GFP moieties on the mother centrioles. Data are presented as Mean±SD. N=3 wing discs, n≥50 centrioles in total for each protein marker. Statistical significance was assessed using an unpaired t test with Welch’s correction (for Gaussian-distributed data) or an unpaired Mann-Whitney test (ns, not significant; ***, P<0.001; ****, P<0.0001). **(C)** The average radial position of each indicated GFP marker – coloured solid line (± 1SD) – is overlaid on a schematic representation of the *Trichonympha*EM-tomogram- derived cartwheel structure (Guichard et al., 2013). The data for Sas-6-GFP and Sas-4-GFP were acquired on the same microscope set-up, but were analysed previously using single-molecule localization microscopy (Gartenmann et al., 2017). They are re-plotted here to indicate the positions of CP110 and Cep97 relative to the outer cartwheel spokes (Sas-6-GFP) and the area linking the cartwheel to the centriole MTs (Sas-4-GFP). CP110 and Cep97 co-localise with the predicted position of the centriole MTs.

Using genetic deletions of *CP110* (Franz et al., 2013) and *Cep97* (Dobbelaere et al., 2020), we found that in early embryos the cytoplasmic levels of each protein was somewhat reduced in the absence of the other, most dramatically for CP110 in the absence of Cep97 (Fig. S1A). Moreover, the centriolar localisation of each protein was largely, although perhaps not completely, dependent on the other (Fig. S1, B and C). These observations support the view that these proteins normally form a complex at the distal end of the centriole MTs (Delgehyr et al., 2012; Dobbelaere et al., 2020; Franz et al., 2013; Kohlmaier et al., 2009; Schmidt et al., 2009; Spektor et al., 2007).

### CP110/Cep97 levels rise and fall on the growing daughter centriole

In the absence of the CP110/Cep97 complex the centriole MTs are dramatically elongated (Dobbelaere et al., 2020; Franz et al., 2013; Schmidt et al., 2009). It is unclear, however, if this is because the MTs on the growing daughter centrioles grow too quickly, or because the centriole MTs on the mother centriole are not properly “capped” to prevent inappropriate MT growth, or both. To determine whether the CP110/Cep97 complex was recruited to growing daughter centrioles, we examined the dynamics of CP110/Cep97 localisation to centrioles in living fly embryos, monitoring their recruitment over nuclear cycles 11-13 (Movies S1 and S2). Unexpectedly, uCP110-GFP and uGFP-Cep97 were recruited to centrioles in a cyclic manner, with centriolar levels being lowest during early mitosis and peaking in approximately mid-S-phase (Fig. 2, A and B). The recruitment dynamics of both proteins appeared to be in phase with each other, and this was confirmed in embryos co-expressing uCP110-GFP and uRFP-Cep97 (Fig. 2C; Movie S3). In contrast, the peak of uRFP-Cep97 recruitment was largely out of phase with the previously described (Aydogan et al., 2020) centriolar oscillation in Plk4-GFP levels, which peak in late-mitosis/early-S-phase (Fig. S2).

**Figure 2.**
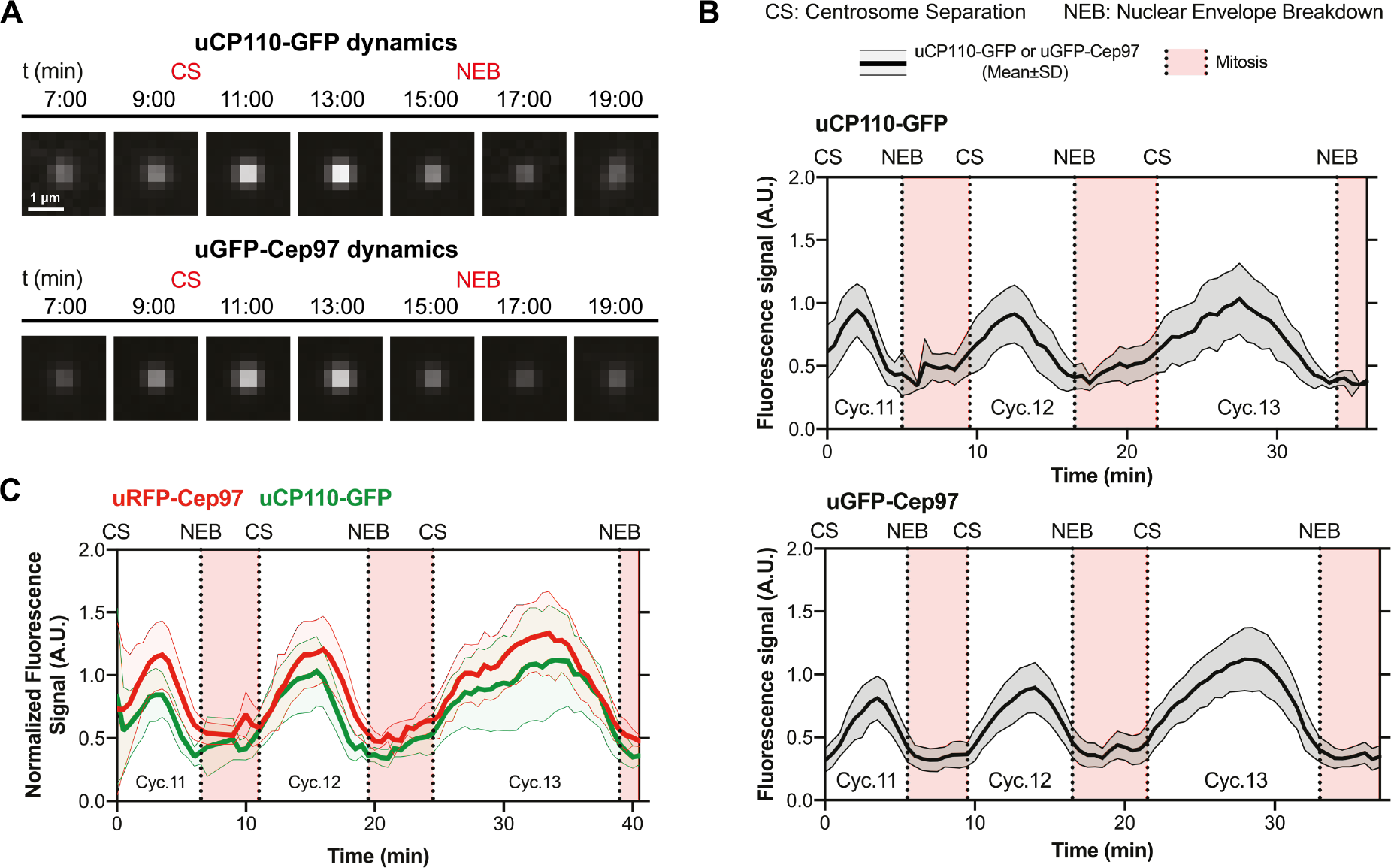
CP110 and Cep97 are recruited to centrioles in a cyclical manner during each nuclear cycle. **(A)** Micrographs from two different embryos illustrate how the centriolar levels of uCP110-GFP and uGFP-Cep97 vary over time during nuclear cycle 12 – obtained by superimposing all the uCP110-GFP (n=150) or uGFP-Cep97 (n=115) centriole foci at each time point. CS=Centrosome Separation, NEB=Nuclear Envelope Breakdown. **(B)** Graphs quantify the centrosomal fluorescence levels (Mean±SD) of uCP110-GFP or uGFP-Cep97 in an individual embryo during cycles 11-13. The graphs are representative examples from 4 independent embryos expressing either uCP110-GFP or uGFP-Cep97 with an average of n=66 or 51 centrioles analysed per embryo, respectively. **(C)** Graph quantifies the centrosomal fluorescence levels (Mean±SD) of uCP110-GFP and uRFP-Cep97 co-expressed in an individual embryo during cycles 11-13. The graph is representative of 6 independent embryos with an average of n=71 centrioles analysed per embryo.

To test whether these dynamics represent changes in CP110/Cep97 levels at the mother and/or at the growing daughter centriole, we examined uCP110- GFP or uGFP-Cep97 recruitment to centrioles in embryos co-expressing Asl-mCherry using Airy-scan super resolution microscopy. Although in our hands 3D-SIM has better resolution than Airy-scan, it is more light intensive and takes longer to acquire images, so we could not use 3D-SIM to follow the dynamic behaviour of these proteins through a nuclear cycle. The Airy-scan analysis revealed that uCP110-GFP and uGFP-Cep97 levels appeared to decrease slightly on mother centrioles as S-phase progressed, while exhibiting a clear cyclical behaviour on the growing daughter (Fig. 3, A and B).

**Figure 3.**
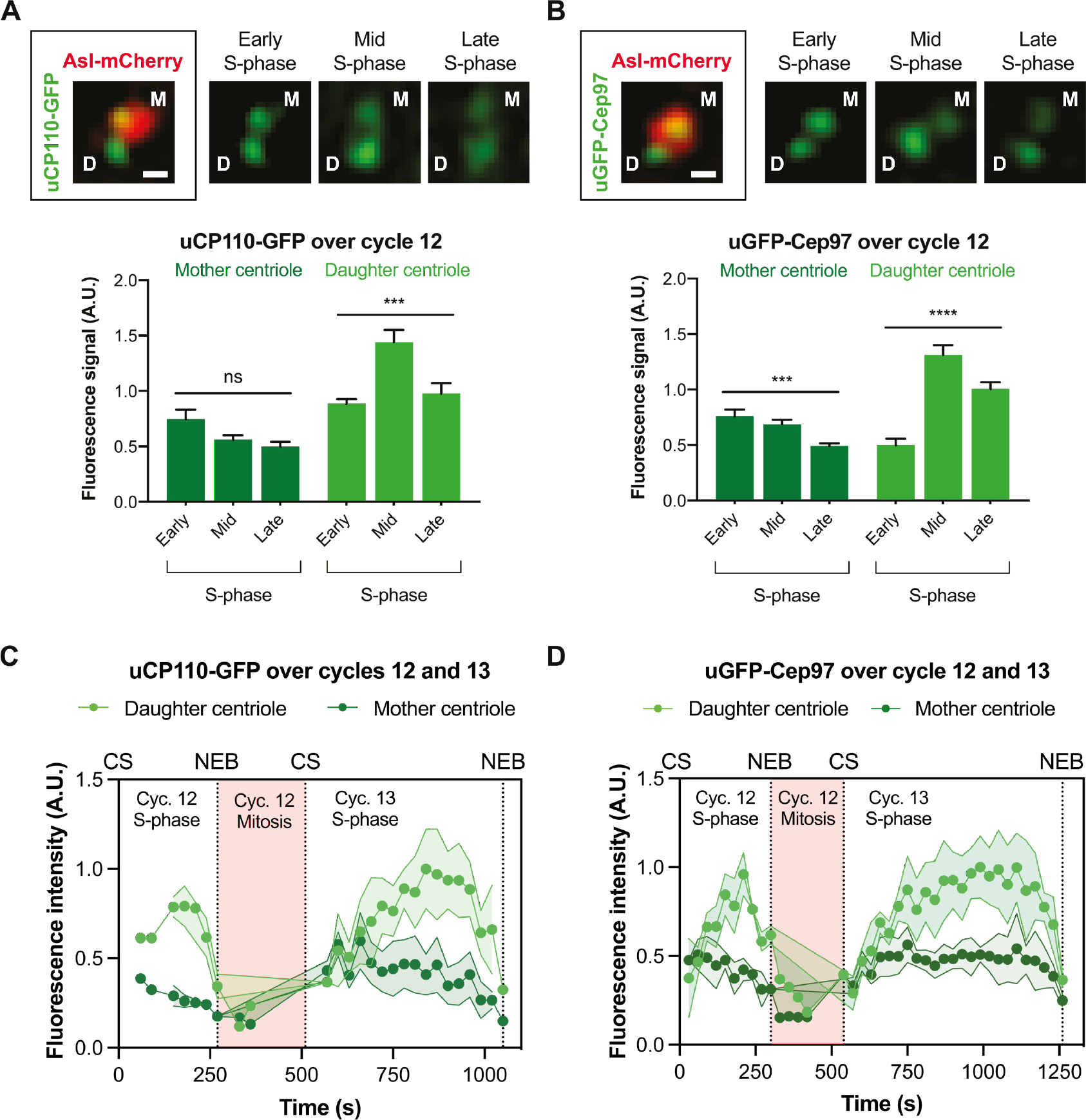
The cyclical recruitment of CP110-GFP and GFP-Cep97 during each nuclear cycle occurs largely at the growing daughter centriole. (A and B) Airy-scan micrographs of centrioles at the indicated stages of S-phase in embryos that express Asl-mCherry and either uCP110-GFP (A) or uGFP-Cep97 (B); D=daughter centriole; M=mother centriole; Scale bars=0.2 μm. Bar charts quantify the centriolar levels (Mean±SEM) of uCP110-GFP (A) or uGFP-Cep97 (B) on the mother (dark green bars) and daughter (light green bars) centrioles at various stages of S-phase. N≥7 embryos. For uCP110- GFP, n=1-9 centrioles per embryo. For uGFP-Cep97, n=1-14 centrioles per embryo. Statistical significance was assessed using an ordinary one-way ANOVA test (for Gaussian-distributed data) or a Kruskal-Wallis test (ns, not significant; ***, P<0.001; ****, P<0.0001). (C and D) Graphs quantify the fluorescence intensity (Mean±SD) acquired using Airy-scan microscopy of uCP110-GFP (C) or uGFP-Cep97 (D) on mother (dark green) and daughter (light green) centrioles in individual embryos over cycles 12-13. CS = Centrosome Separation, NEB = Nuclear Envelope Breakdown. For uCP110-GFP, n=1-3 or 1-8 daughter and mother centrioles per time point in cycles 12 and 13, respectively. For uGFP-Cep97, n=1-4 or 1-10 daughter and mother centrioles per time point in cycles 12 and 13, respectively. Note that the numbers of centrioles analysed in these experiments is relatively low because the Airy-scan system has a small field of view (so we can track fewer centrioles), and because centrioles have to be unambiguously assigned as mothers or daughters. This was not always possible, and was particularly challenging during mitosis when the centrioles move rapidly within the embryo, and when the daughter centrioles are also starting to load Asl-mCherry as they mature into new mothers.

To examine whether the gradual decrease in signal on the mother centriole was due to photobleaching, we attempted to use the Airy-scan to follow mother and daughter centrioles as they finished S-phase of cycle 12 and then proceeded through mitosis and S-phase of nuclear cycle 13. This proved challenging: centrioles are very mobile and very dim during mitosis (making them hard to track), and daughter centrioles start to load Asl-mCherry during mitosis as they mature into mothers (Novak et al., 2014), making it difficult to unambiguously distinguish the two centrioles in a separating pair.

Nevertheless, although only a small number of centrioles could be unambiguously tracked and assigned during mitosis, it appeared that CP110/Cep97 levels on the mother centrioles increased as the embryos progressed from mitosis into the next S-phase (Fig. 3, C and D). Thus, CP110 and Cep97 levels probably rise and fall at both centrioles during each nuclear cycle, but this is more pronounced on growing daughters.

These observations are in contrast with a recent report in syncytial fly embryos that Cep97-GFP expressed under its endogenous promoter is first recruited to new-born centrioles late in S-phase, after they have already fully elongated (Dobbelaere et al., 2020). We wondered, therefore, whether our observations might be an artefact of overexpression, as the proteins we used here are overexpressed from the *Ubq*-promoter by ∼2-5X compared to their endogenous proteins (Fig. S1A; Fig. 4A and B). We therefore analysed the recruitment of CP110-GFP and Cep97-GFP to centrioles during nuclear cycle 12 using transgenic lines in which each protein was expressed from their endogenous promoters (eCP110-GFP and eCep97-GFP – the same line used by Dobbelaere et al.). Interestingly, both fusion proteins were expressed at ∼2-5X lower levels than their endogenous proteins (Fig. 4, A and B), but they exhibited a clear cyclical recruitment during S-phase (Fig. 4, C and D). Thus, in our hands, CP110-GFP and GFP-Cep97 are recruited to centrioles in a cyclical manner if they are either moderately overexpressed (from the *Ubq*- promoter) or moderately under-expressed (from their endogenous promoters), strongly arguing that this behaviour is not an artefact of over- or under- expression.

**Figure 4.**
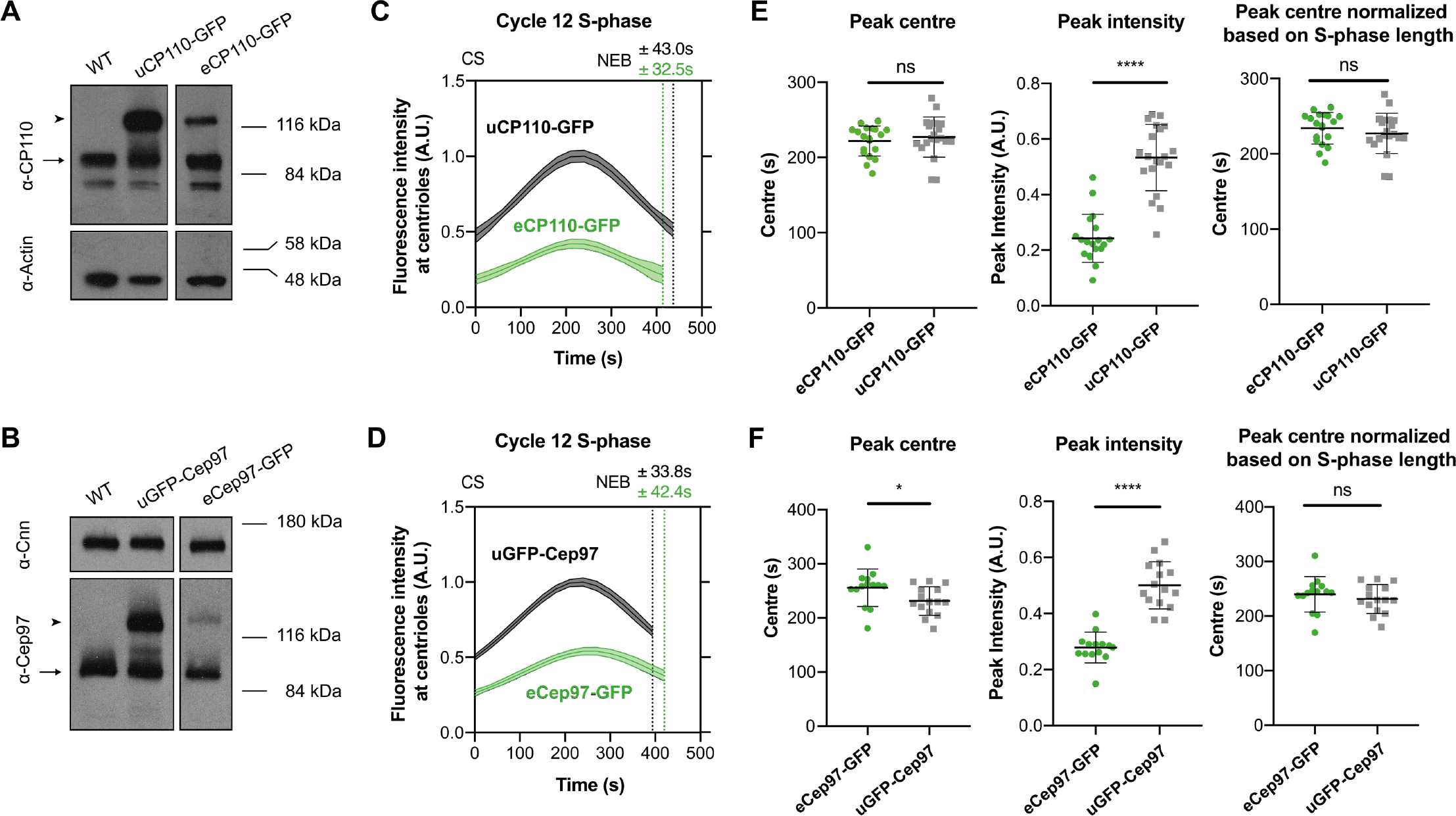
CP110/Cep97 are recruited to centrioles faster and to higher levels when they are expressed at higher levels. **(A and B)** Western blots comparing CP110 or Cep97 expression levels in WT embryos or embryos expressing one copy of either eCP110-GFP or uCP110- GFP (A), or eCep97-GFP or uGFP-Cep97 (B). Endogenous CP110 and Cep97 are indicated with arrows, the GFP-tagged proteins with arrowheads. Actin and Cnn are shown as loading controls. **(C and D)** Graphs compare how the levels (Mean±SEM) of centriolar CP110/Cep97-GFP change during nuclear cycle 12 in embryos expressing (C) uCP110-GFP or eCP110-GFP and (D) uGFP-Cep97 or eCep97-GFP, as indicated. CS=Centrosome Separation, NEB=Nuclear Envelope Breakdown. **(E and F)** Bar charts quantify several parameters (Mean±SD) of the recruitment dynamics derived from the profiles shown in (C and D). N≥14 embryos per group, n≥9 centrioles per embryo. Statistical significance was assessed using an unpaired t test with Welch’s correction (for Gaussian-distributed data) or an unpaired Mann-Whitney test (ns, not significant; *<0.05; ****, P<0.0001).

### The growth period of the centriole MTs does not appear to be set by CP110/Cep97 recruitment dynamics

The centriolar CP110/Cep97 levels peaked in approximately mid-S-phase (Fig. 4, C and D), which is about the same time that the daughter centrioles appear to normally stop growing (Aydogan et al., 2018). We wondered, therefore, whether the gradual accumulation of CP110/Cep97 at the distal end of the growing daughter MTs might influence the time at which the centriole MTs stop growing. We noticed, however, that although significantly more CP110-GFP or GFP-Cep97 was recruited to centrioles when the proteins were overexpressed from the *Ubq*-promotor than when expressed from their endogenous promotors (judged by their *peak intensity*), the relative phase of recruitment (judged by their absolute and S-phase normalized *peak centres*) was hardly altered (Fig. 4, E and F). Indeed, inspection of the recruitment dynamics revealed that the centriolar levels of uCP110-GFP and uGFP- Cep97 were already similar or higher at the start of S-phase than the peak levels attained in the eCP110-GFP and eGFP-Cep97 embryos (Fig. 4, C and D). Thus, it seems unlikely that the centriole MTs stop growing when CP110/Cep97 levels reach a critical threshold-level at the distal end, as the centriole MTs would hardly grow at all in S-phase in the uCP110-GFP and uGFP-Cep97 embryos if this were the case. Although we cannot directly measure the growth of the centriole MTs, we have shown previously that overexpressing CP110-GFP from the *Ubq*-promoter only modestly shortens centriole-MT length in wing disc cells (Franz et al., 2013).

### CP110/Cep97 recruitment dynamics appear to be entrained by the Cdk/Cyclin cell cycle oscillator

These overexpression studies also suggest that centriolar CP110/Cep97 levels do not normally peak in mid-S-phase because the centriolar binding sites for the complex are saturated – as significantly more CP110/Cep97 can be recruited to centrioles when these proteins are overexpressed (Fig. 4). We wondered, therefore whether the CP110/Cep97 recruitment dynamics might be regulated by the core Cdk/Cyclin cell cycle oscillator (CCO) that drives the nuclear divisions in these early embryos. During nuclear cycles 11-13, the rate of CCO activation during S-phase gradually slows, leading to the lengthening of S-phase at successive cycles (Farrell and O’Farrell, 2014; Liu et al., 2021). Interestingly, there was a strong correlation between the timing of the CP110/Cep97 peak and S-phase length for both uCP110-GFP and uGFP-Cep97 at all nuclear cycles (average r=0.86±0.19; p<0.0001 except for uGFP-Cep97 in Cycle 11, where p=0.04) (Fig. 5, A and B). This correlation was also observed in the eCP110-GFP and eCep97-GFP lines that we analysed during nuclear cycle 12 (average r=0.86±0.08; p=<0.0001) (Fig. 5, C and D). These observations suggest that the recruitment of CP110/Cep97 to centrioles might be regulated by the CCO.

**Figure 5.**
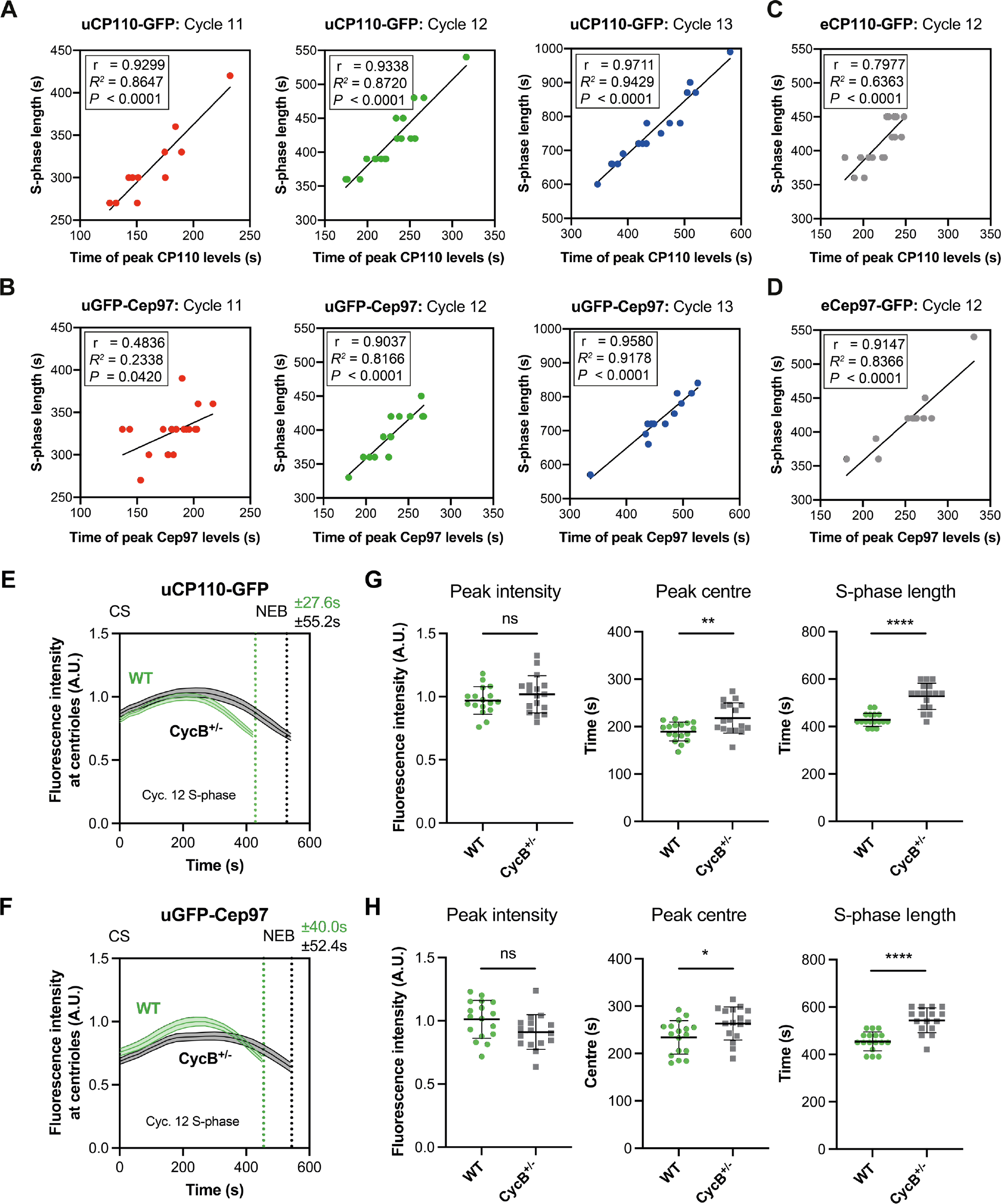
The phase of CP110/Cep97 recruitment is strongly correlated to, and regulated by, the progression of the cell cycle in fly embryos. Scatter plots show the positive correlation between S-phase length and the peak time of centriolar uCP110-GFP (A) and uGFP-Cep97 (B) levels in cycle 11–13, and of eCP110-GFP (C) and eCep97-GFP (D) levels in cycle 12. The plots were regressed using the line function in GraphPad Prism 8. Correlation strength was examined using Pearson’s correlation coefficient (0.40<r<0.60 = moderate; r>0.60 = strong), and the statistical significance of the correlation was determined by the p-value. (E and F) Graphs compare how the levels (Mean±SEM) of centriolar CP110/Cep97-GFP change during nuclear cycle 12 in embryos expressing uCP110-GFP (E) and uGFP-Cep97 (F) in WT or CycB^+/-^ embryos, as indicated. CS=Centrosome Separation, NEB=Nuclear Envelope Breakdown. (G and H) Bar charts quantify several parameters (Mean±SD) of the recruitment dynamics derived from the profiles shown in (E and F respectively). N≥16 embryos per group, n≥50 centrioles per embryo. Statistical significance was assessed using an unpaired t test with Welch’s correction (ns, not significant; *<0.05; **<0.01; ****, P<0.0001).

To test if this was the case, we examined the kinetics of uCP110-GFP and uGFP-Cep97 recruitment in embryos in which we halved the genetic dose of Cyclin B. This perturbation extends S-phase length (presumably because it takes more time for Cdk/Cyclin levels to reach the threshold required to trigger mitotic entry) and it also extended the period of CP110/Cep97 recruitment (as measured by the time taken to reach peak intensity) (Fig. 5, E–H). This is consistent with the idea that as S-phase progresses, gradually increasing Cdk/Cyclin activity (Deneke et al., 2016) is either directly or indirectly responsible for switching off the ability of the centrioles to recruit CP110/Cep97.

### CP110/Cep97 influence the rate and period of centriole cartwheel growth

We next tested whether CP110/Cep97 might instead influence the growth of the central cartwheel. This may seem counterintuitive, as we previously showed that the daughter centriole cartwheel grows preferentially from the proximal end (Aydogan et al., 2018), whereas CP110/Cep97 are recruited to the distal end of the growing daughter. Nevertheless, it is striking that the ability of Plk4 to promote centriole overduplication in human cells requires CP110 (Kleylein-Sohn et al., 2007) and the phosphorylation of CP110 by Plk4 is required for efficient centriole assembly in at least some systems (Lee et al., 2017), hinting at possible crosstalk between Plk4 and the CP110/Cep97 complex.

We previously used the incorporation of the core centriole cartwheel component Sas-6-GFP as a proxy to measure centriole cartwheel growth (Aydogan et al., 2018), so we applied this assay to determine the parameters of cartwheel growth in embryos that either lacked or overexpressed CP110 or Cep97 (Fig. S1A). Embryos with these mutations are viable and display no obvious developmental defects (Fig. S1D). As a control, we confirmed that Sas-6-GFP was preferentially incorporated only into the proximal end of the growing daughter centrioles even in the absence of CP110 or Cep97 (Fig. S3). This excludes the possibility that Sas-6 can inappropriately incorporate into the distal-end of either the mother centrioles or the growing daughter centrioles if they are not capped by CP110 or Cep97.

To our surprise, the centriole cartwheel grew more quickly when CP110 or Cep97 were absent, and more slowly when either protein was overexpressed (Fig. 6), demonstrating that CP110/Cep97 levels in the embryo influence the rate of cartwheel growth. Cartwheel growth is normally regulated homeostatically: when cartwheels grow faster, they tend to grow for a shorter period, and *vice versa* (Aydogan et al., 2018). This homeostatic regulation appeared to be largely, but not perfectly, maintained in embryos in which the levels of CP110 or Cep97 were altered: the cartwheels grew faster, but for a shorter period in embryos lacking CP110 or Cep97, and more slowly, but for a longer period in embryos overexpressing CP110 or Cep97 (Fig. 6). The changes in the growth period, however, were not sufficient to compensate for the more dramatic changes in the growth rate, so the cartwheels were slightly longer in embryos lacking CP110 or Cep97, and slightly shorter in embryos overexpressing CP110 or Cep97 (Fig. 6). This is in good agreement with our previous electron microscopy studies showing that the core centriole structure in wing disc cells lacking CP110 is slightly longer (although the centriole MTs are dramatically elongated), while in cells overexpressing CP110 the centrioles are slightly shorter (Franz et al., 2013).

**Figure 6.**
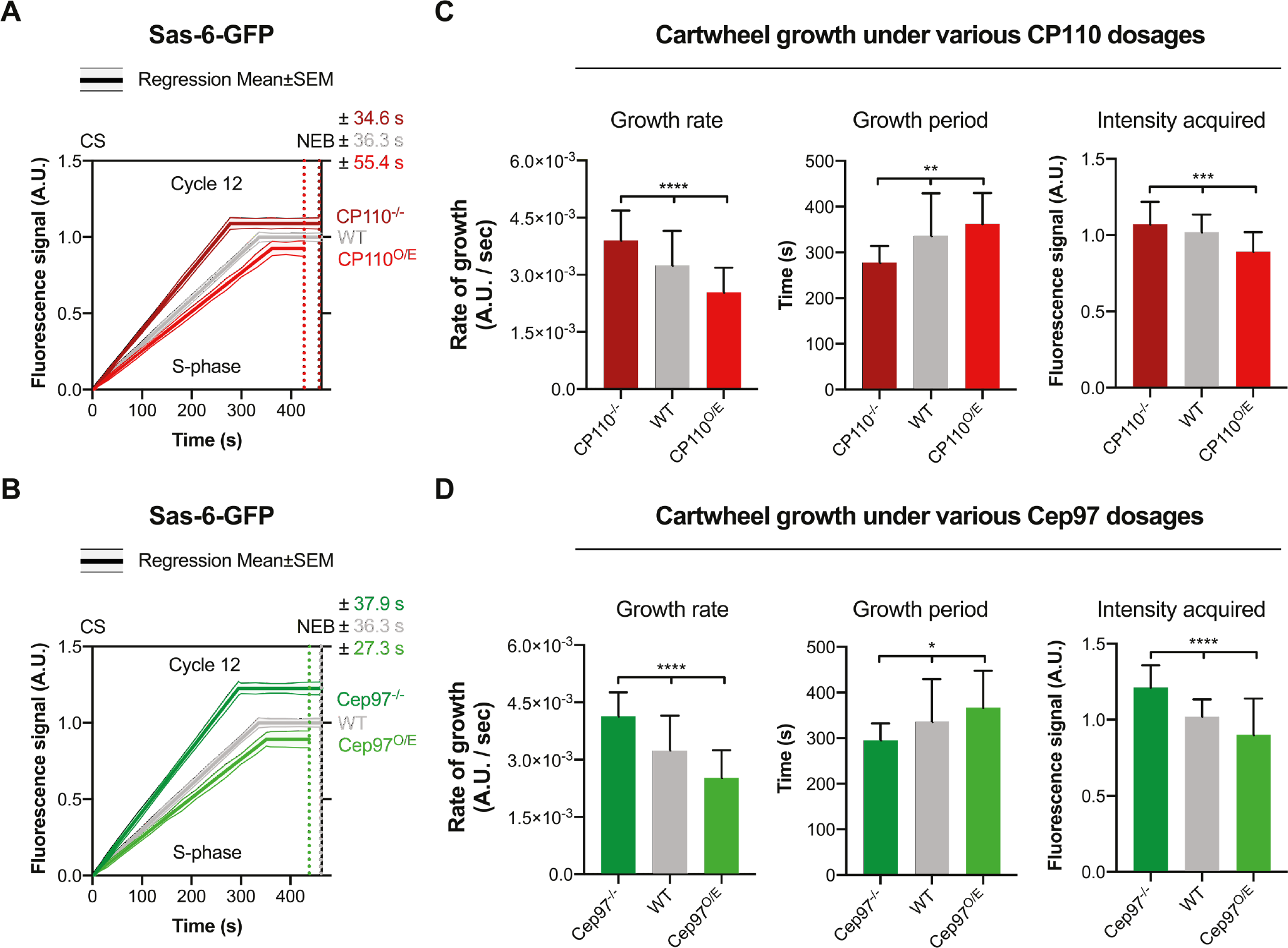
CP110 and Cep97 levels influence the rate and period of centriole cartwheel growth. **(A and B)** Graphs compare the Sas-6-GFP incorporation profile (Mean±SEM) – as a proxy for centriole cartwheel growth (Aydogan et al., 2018) – during nuclear cycle 12 in WT embryos or in embryos lacking or overexpressing CP110 (A) or Cep97 (B). **(C and D)** Bar charts quantify several parameters of cartwheel growth (Mean±SD) derived from the profiles shown in (A) and (B) respectively. N≥14 embryos per group, n≥40 centrioles on average per embryo. Statistical significance was assessed using an ordinary one-way ANOVA test (for Gaussian-distributed data) or a Kruskal-Wallis test (*, P<0.05; **, P<0.01; ***, P<0.001; ****, P<0.0001).

### CP110/Cep97 levels influence the Plk4 oscillation at the proximal end of the growing daughter centriole

To test whether the CP110/Cep97 complex influences cartwheel growth by altering the centriolar Plk4 oscillation, we examined the Plk4-GFP oscillations in nuclear cycle 12 in embryos where CP110 or Cep97 were either absent or were individually overexpressed. These perturbations altered both the amplitude and phase of the Plk4 oscillation at the proximal end of the growing daughter (Fig. 7).

**Figure 7.**
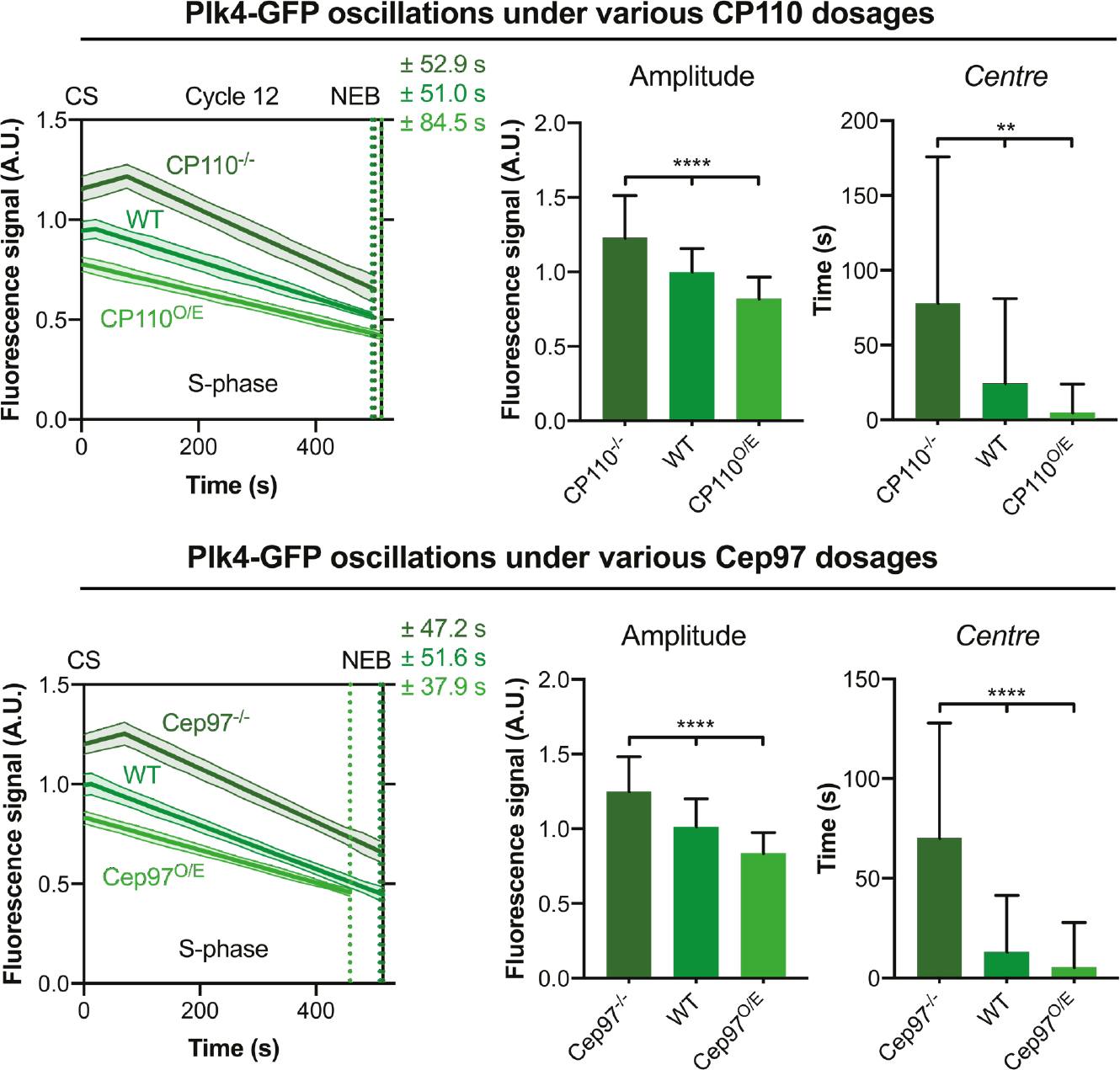
CP110 and Cep97 levels influence the parameters of the Plk4 oscillation. Graphs show the fitted oscillation in centriolar Plk4-GFP levels (Mean±SEM) in S-phase of nuclear cycle 12 in embryos expressing various levels of either CP110 or Cep97. The Plk4 oscillation was previously shown to influence the parameters of centriole growth (Aydogan et al., 2020). Corresponding bar charts compare the amplitude and centre (Mean±SD) of the fitted Plk4-GFP oscillation under the indicated conditions. N≥16 embryos per group, n≥45 centriole pairs per embryo. Statistical significance was assessed using an ordinary one-way ANOVA test (for Gaussian-distributed data) or a Kruskal-Wallis test (**, P<0.01; ****, P<0.0001).

We previously showed that in WT embryos the Plk4 oscillation exhibits adaptive behaviour: as its amplitude tends to decrease from nuclear cycle 11 to 13, so its period tends to increase -- and we hypothesised that the progressively decreasing amplitude helps to slow the growth rate at successive cycles, while the progressively increasing period helps to increase the cartwheel growth period at successive cycles (Aydogan et al., 2020).

Strikingly, the lack of CP110 or Cep97 led to an increase in the amplitude of the Plk4 oscillation – consistent with the faster rate of cartwheel growth we observe (Fig. 6) – but the peak of the Plk4 oscillation was shifted to later in S-phase, even though the cartwheels in these embryos grow for a shorter period. Similarly, while the overexpression of CP110 or Cep97 led to a decrease in the amplitude of the Plk4 oscillation – consistent with the slower rate of cartwheel growth we observe (Fig. 6) – the peak of the Plk4 oscillation was shifted to earlier in S-phase, even though the cartwheels in these embryos grow for a longer period (see Discussion).

### Altering CP110/Cep97 levels does not detectably alter the cytoplasmic levels of Plk4, or *vice versa*, in embryos

We wondered whether CP110/Cep97 might influence Plk4 abundance in the cytoplasm. As described previously, in our hands the cytosolic concentration of Plk4-monomeric NeonGreen expressed from its own promoter was too low to be measured by conventional *Fluorescence Correlation Spectroscopy* (FCS), so we used *Peak Counting Spectroscopy* (PeCoS) instead (Aydogan et al., 2020). Varying the cytoplasmic dosage of CP110 or Cep97 did not detectably alter cytoplasmic Plk4 levels (Fig. 8A). In addition, we used FCS and western blotting to measure the cytoplasmic concentration or total amount, respectively, of uCP110-GFP and uGFP-Cep97 in *Plk4^1/2^* embryos or in embryos carrying a previously described mutated form of Plk4 with reduced kinase activity (*Plk4^RKA^*) (Aydogan et al., 2018). Neither the levels nor the kinase activity of Plk4 appeared to influence CP110 and Cep97 levels (Fig. 8, B and C). Thus, any crosstalk between CP110/Cep97 and Plk4 does not appear to rely on their ability to influence each other’s cytoplasmic abundance in embryos.

**Figure 8.**
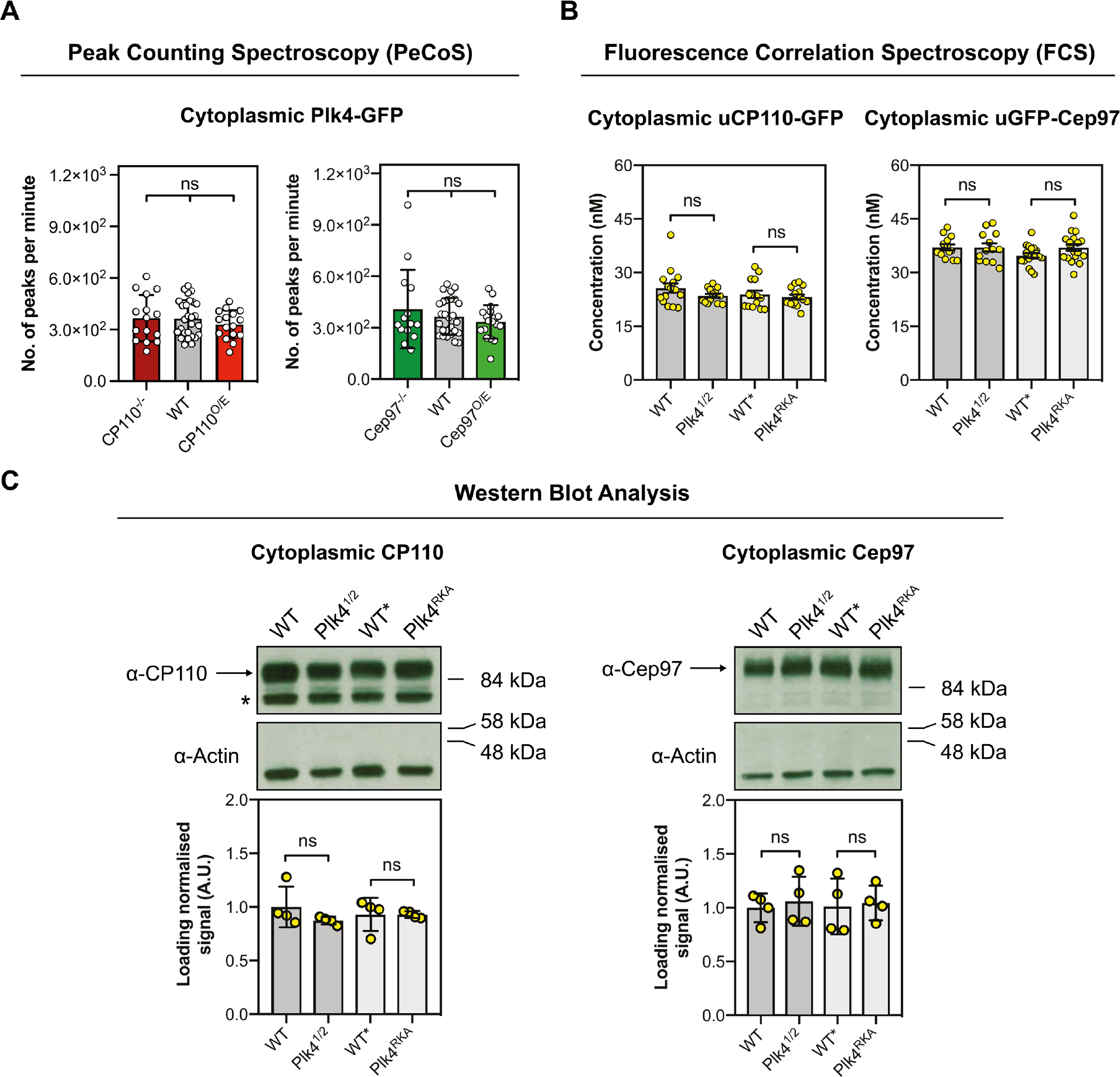
Altering the cytoplasmic levels or activity of Plk4 does not detectably alter the cytoplasmic levels of CP110 or Cep97 and vice versa. **(A)** Bar charts quantify PeCoS measurements (Mean±SD) of Plk4-GFP in embryos expressing various levels of CP110 and Cep97. Every data point represents 1x 180 sec measurement from an individual embryo. Statistical significance was assessed using an ordinary one-way ANOVA test (for Gaussian-distributed data) or a Kruskal-Wallis test (ns, not significant). **(B)** Bar charts quantify the background-corrected FCS measurements (Mean±SD) of uCP110-GFP or uGFP-Cep97 under the indicated conditions. Each data point represents the average of 4-6 recordings from each embryo measured. Statistical significance was assessed using an ordinary unpaired t-test (for Gaussian-distributed data) or a Mann-Whitney test (ns, not significant). **(C)** (Upper panel) Western blots show the cytoplasmic expression of endogenous CP110 and Cep97 (arrows) under the same conditions used to measure the concentration of uCP110-GFP and uGFP-Cep97 by FCS in (B). Actin is shown as a loading control, and prominent non-specific bands are indicated (*). Representative blots are shown from four technical repeats. (Lower panel) Bar charts quantify the loading-normalised levels (Mean±SD) of CP110 and Cep97 from the four technical repeats. Statistical significance was assessed using a Mann-Whitney test (ns, not significant).

## Discussion

Here we show that CP110 and Cep97 are recruited to the distal-end of daughter centriole MTs in a cyclical manner as they grow during S-phase, with levels peaking, and then starting to decline at ∼mid-S-phase, which is normally when the centrioles appear to stop growing in these embryos (Aydogan et al., 2018). These recruitment dynamics, however, do not appear to play a major part in determining the period of daughter centriole MT growth, and our findings strongly suggest that centriole MTs do not stop growing when a threshold level of CP110/Cep97 accumulates at the centriole distal end.

Thus, although in many systems the centriole MTs are dramatically elongated in the absence of CP110 or Cep97 (Dobbelaere et al., 2020; Franz et al., 2013; Kohlmaier et al., 2009; Schmidt et al., 2009; Sharma et al., 2021; Winey and O’Toole, 2014), we speculate that this is largely due to a failure to properly “cap” these MTs after they have finished growing, rather than because the centriole MTs grow too quickly as the new daughter centriole is being assembled.

CP110 and Cep97 levels do not peak at centrioles because the proteins reach saturating levels on the centriole MTs, as the amount of CP110/Cep97 recruited to centrioles is increased when either protein is overexpressed. It is unclear how these proteins interact specifically with the distal-ends of the centriole MTs, but we conclude that their binding sites are normally far from saturated, at least in the rapidly cycling *Drosophila* embryo. Importantly, the phase of CP110/Cep97 recruitment is influenced by the activity of the core Cdk/Cyclin cell cycle oscillator (CCO). We suspect, therefore, that the cyclical recruitment dynamics of CP110/Cep97 in these embryos may simply reflect the ability of these proteins to bind to centrioles when Cdk/Cyclin activity is low, but not when it is high. CP110 was originally identified as a CDK substrate (Chen et al., 2002), and presumably the CCO modifies (perhaps by phosphorylating) CP110 and/or Cep97 and/or their centriolar recruiting factor(s) to inhibit recruitment as cells prepare to enter mitosis. It is presently unclear why it might be important to prevent CP110/Cep97 binding to centrioles during mitosis.

Perhaps surprisingly, we show that CP110/Cep97 levels can influence the growth of the centriole cartwheel, at least in part, by altering the parameters of the Plk4 oscillation at the base of the growing daughter centrioles. This reveals an unexpected crosstalk between proteins that are usually thought to influence events at the proximal end of the cartwheel (Plk4) and at the distal end of the centriole MTs (CP110/Cep97). We currently do not understand how CP110/Cep97 might influence Plk4’s behaviour, but our data suggests they do not alter each other’s abundance in the cytoplasm. Nevertheless, it may be that Plk4 and CP110/Cep97 interact in the cytoplasm, and this interaction influences the amount of Plk4 available for recruitment to the centriole (explaining why less Plk4 is recruited when these proteins are overexpressed and more is recruited with they are absent). Alternatively, perhaps these proteins interact at the centriole during the very early stages of daughter centriole assembly, when they are all present at the nascent site of assembly but have not yet been spatially separated by the growth of the daughter centriole. Clearly it will be important to test whether Plk4 and CP110/Cep97 interact in *Drosophila* embryos and, if so, how this interaction is regulated in space and time.

CP110 and Cep97 are not essential for centriole duplication in mice or flies (Dobbelaere et al., 2020; Franz et al., 2013; Yadav et al., 2016), but CP110 is required for Plk4-induced centriole overduplication in cultured human cells (Kleylein-Sohn et al., 2007), and Plk4 can interact with and phosphorylate CP110 to promote centriole duplication in these cells (Lee et al., 2017). Thus, although the physiological significance and molecular mechanism of Plk4 and CP110/Cep97 crosstalk is currently unclear, this crosstalk may be conserved in other species.

Finally, it is important to note that changing the levels of CP110/Cep97 influences the Plk4 oscillation in a surprising way. In the absence of CP110/Cep97, the cartwheel grows faster and for shorter period, but the Plk4 oscillation has a higher amplitude, and a longer period. Our previous observations would suggest that faster centriole growth for a shorter period would be associated with Plk4 oscillation that has a higher amplitude, but a *shorter* period (Aydogan et al., 2020). One way to potentially explain this conundrum is if Plk4 is more active in the absence of CP110/Cep97 – so the cartwheel would be built faster, but for a shorter period (Aydogan et al., 2020), as we observe – but the inactivated Plk4 is not efficiently released from its centriolar receptors (so Plk4 would accumulate at centrioles to a higher level and for a longer period). Clearly further work is required to understand how the Plk4 oscillation drives cartwheel assembly, and how this process is influenced by CP110/Cep97.

## Acknowledgements

We thank Jeroen Dobbelaere and Alex Dammermann for sharing the eGFP- Cep97 line and anti-Cep97 antibodies prior to publication, and members of the Aydogan and Raff laboratories for critically reading the manuscript. Super-resolution microscopy was carried out at the Micron Oxford Advanced Bioimaging Unit, funded by a Strategic Award from the Wellcome Trust (107457). The research was funded by a Wellcome Trust Senior Investigator Award (104575 and 215523; T.L.S., M. M., S. S., A. W. and J.W.R.), an Edward Penley Abraham Scholarship (M.G.A.), a Sandler Foundation Investigator Award (7029760; M.G.A.), a UCSF PBBR New Frontiers Research Award (2017078; M.G.A.), a Wellcome Trust PhD Studentship (203855; L.E.H.), and a Ludwig Institute for Cancer Research funding (F.Y.Z.).

## Author contributions

This study was conceptualised by M.G.A., L.E.H. and J.W.R. Investigation was done by M.G.A., L.E.H., T.L.S., M.M., S.S. and A.W. Data were analysed by M.G.A., L.E.H., T.L.S. and A.W. Methodology was developed by M.G.A., L.E.H. and J.W.R. Project was administrated by M.G.A., L.E.H. and J.W.R. Resources were shared/made by M.G.A., L.E.H., T.L.S., M.M., S.S., S.S.W., X.L. and F.Y.Z. Software work was carried out by M.G.A., L.E.H. and F.Y.Z. Overall supervision was done by M.G.A. and J.W.R. Validation experiments/analyses were carried out by M.G.A., L.E.H., T.L.S and S.S. Finally, M.G.A., L.E.H. and J.W.R. wrote, reviewed and edited the manuscript with input from all authors.

## Data, material and code availability

The script to automatize the CP110- and Cep97-GFP ring measurements analysis (of the super-resolution microscopy data) is available in the following link: <https://github.com/RaffLab/SIM-centriole-ring-measurement>. The rest of the data is available upon request.

## Materials and Methods

### Drosophila melanogaster stocks and husbandry

*D. melanogaster* stocks used in this study are listed in Supplementary Table 1, and the lines generated and tested here are listed in Supplementary Table 2. To generate the Ubq-CP110 (uCP110) construct, a stop codon was introduced into a previously generated pDONR-CP110L (containing a full length *CP110* cDNA) (Franz et al., 2013) by site directed mutagenesis using Quikchange II XL mutagenesis kit (Agilent technologies). This was then recombined with the pUbq-empty vector (This paper; details available on request). To generate the Ubq-Cep97 (uCep97) construct, the pZeo-CG3980- NT vector (Dobbelaere et al., 2008) was directly recombined with the pUbq- empty vector.

The pDONR-Zeo-eCP110 cDNA construct was cloned by assembling two fragments: the ∼2kb region upstream of the cp110 start codon and the pDONR-Zeo-CP110 cDNA vector containing the long isoform of CP110 minus the stop codon (Franz et al., 2013), using NEBuilder HiFi assembly (NEB).

The cDNA construct pDONR-Zeo-eCP110 was then recombined with an mGFP-CT empty destination vector (pNoP-mGFP-CT-DEST; This paper – details available upon request) via Gateway Technology (Thermo Fisher Scientific). The pNoP-mGFP-CT-DEST was made by removing the Ubiquitin promoter from a previously published (Basto et al., 2008) Ubq-mGFP-CT destination vector.

Primer sequences used to introduce a stop codon for the uCP110 construct, to amplify the *cp110* promoter region, to amplify the pDONR-Zeo vector containing the *cp110* sequence, and to clone the C-terminal fragment aa 329- 807 of Cep97 into the pDONR vector are listed in Supplementary Table 3.

Transgenic flies were generated by the Fly Facility in the Department of Genetics, University of Cambridge (UK).

Flies were maintained at 18°C or 25°C on *Drosophila* culture medium (0.77% agar, 6.9% maize, 0.8% soya, 1.4% yeast, 6.9% malt, 1.9% molasses, 0.5% propionic acid, 0.03% ortho-phosphoric acid and 0.3% nipagin) in vials or in bottles. For embryo collections, 25% cranberry-raspberry juice plates (2% sucrose and 1.8% agar with a drop of yeast suspension) were used.

### Embryo collections

For all imaging experiments, embryos were collected for 1 h at 25°C, and then aged for 45 min to 1 h. Before imaging, embryos were dechorionated by hand, mounted on a strip of glue painted on a 35 mm glass bottom petri dish with 14 mm micro-well (MatTek) and desiccated for 1 min at 25°C. Embryos were then covered with Voltalef^®^ oil (ARKEMA).

### Hatching experiments

In order to measure embryo hatching rates, 0-3 h embryos were collected and aged for 24 h, and the % of embryos that hatched out of their chorion was scored. 5 technical repeats were carried out over multiple days, and at least 120 embryos were analysed for each genotype per repeat.

### Image acquisition, processing and analysis

#### Airy-scan super resolution microscopy

Living embryos were imaged using an inverted Zeiss 880 microscope fitted with an airy-scan detector. The system was equipped with Plan-Apochromat 63x/1.4 NA oil lens. 488 nm argon and 561 nm diode lasers were used to excite GFP and RFP (or mCherry) respectively. Stacks of 5 slices at 0.2 μm intervals were collected with a zoom value of 24.41 pixels/μm. Focus was re- adjusted in between image collection. Images were airy-processed in 3D with a strength value of *Auto* (∼6) or 6.5.

Fluorescence Recovery After Photobleaching (FRAP) experiments were performed on the same Zeiss 880 system. Settings for the sequence of events needed for FRAP experiments were as follows: **(1)** Acquisition of a single Z-stack in Airy-scan mode (*Pre-bleach* in Fig. S2A); **(2)** multi-spot serial photo-bleaching (4 regions at a time); **(3)** acquisition of the photo-bleached image in Airy-scan mode (*Bleach* in Fig. S3A); **(4)** acquisition of the post-bleach images in Airy-scan mode (*Post-bleach* in Fig. S3A).

This protocol was also used to examine the site where Sas-6-GFP incorporates into centrioles in WT, CP110^-/-^ and Cep97^-/-^ embryos (Fig. S3B). For this analysis, embryos exiting mitosis of nuclear Cycle 13 were identified and daughter centrioles were allowed to grow for 6 minutes into Cycle 14 – allowing centrioles to grow to approximately half of their final size (Aydogan et al., 2018). The site of new Sas-6-GFP recruitment was then determined using a previously described pipeline (Aydogan et al., 2018). This analysis was performed blind.

#### Spinning Disk Confocal Microscopy

Living embryos were imaged using either a Perkin Elmer ERS Spinning Disk confocal system on a Zeiss Axiovert 200M microscope equipped with Plan-Apochromat 63x/1.4 NA oil DIC lens, or an EM-CCD Andor iXon+ camera on a Nikon Eclipse TE200-E microscope with a Plan-Apochromat 60x/1.42 NA oil DIC lens. 488 and 561 nm lasers were used to excite GFP and RFP (or mCherry) respectively. Confocal sections of 13 (Perkin Elmer) or 17 (Andor) slices with 0.5 μm thick intervals were collected every 30 sec.

Post-acquisition image processing (including image projection, photo- bleaching correction and background subtraction) was carried out using Fiji (NIH, US) as described previously (Aydogan et al., 2018). uCP110-GFP, eCP110-GFP, uGFP-Cep97, eCep97-GFP, Plk4-GFP, uRFP-Cep97 and Sas-6-GFP foci were tracked using TrackMate (Tinevez et al., 2017), a plug-in of Fiji, with track spot diameter size of 1.1 μm. The regressions for Plk4-GFP oscillations and Sas-6-GFP cartwheel growth curves were calculated in Prism 8 (GraphPad Software), as described previously (Aydogan et al., 2018). The regressions for uGFP-Cep97 and eCep97-GFP (or, uCP110-GFP and eCP110-GFP) dynamics were calculated using the *nonlinear regression* (curve fit) function in Prism 8. Discrete curves in S-phase were first fitted against four different functions to decide the best regression model: 1) Lorentzian, 2) Gaussian, 3) Increase – Constant – Decrease, and 4) Increase

– Decrease. Among these models, Lorentzian best fit the data (data not shown). Thus, all the fluorescently tagged Cep97 and CP110 curves in S-phase were regressed with the Lorentzian function. The Lorentzian and Gaussian functions are integral to Prism 8, while the latter two functions were described previously (Alvarez-Rodrigo et al., 2019). In order to plot the dynamics of uCP110-GFP and uRFP-Cep97 together, the highest mean fluorescence signal for each tag in nuclear cycle 12 was scaled to 1 A.U., and this scaling factor was accordingly applied across all cycles.

As previously described (Aydogan et al., 2018), the beginning of S-phase was taken as the time of centrosome separation (“CS”). Entry into mitosis was taken as the time of nuclear envelope breakdown (NEB).

#### 3D-Structured Illumination Microscopy (3D-SIM)

Living embryos were imaged at 21°C using a DeltaVision OMX V3 Blaze microscope (GE Healthcare Life Sciences, UK). The system was equipped with a 60x/1.42 NA oil UPlanSApo objective (Olympus Corp.), 488 nm and 593 nm diode lasers and Edge 5.5 sCMOS cameras (PCO). All image acquisition and post-processing was performed as described previously (Aydogan et al., 2018, 2020). In order to investigate where CP110 and Cep97 radially localizes on the centriole, wing disc preparations were made as previously described (Gartenmann et al., 2017) and imaged with an OMX V2 microscope (GE Healthcare Life Sciences, UK) with a 100x/1.4 NA oil objective (UPlanSApo, Olympus), 488nm diode (Sapphire 488-200, Coherent) and 592nm (F-04306-01, MPB Communications) fiber lasers and Evolve 512 Delta C EMCCD cameras (Photometrics). The orientation of the centrioles and the radial localisation of the various CP110 and Cep97 constructs was assessed following a previously described protocol (Gartenmann et al., 2017) and using our own python code (available at: https://github.com/RaffLab/SIM-centriole-ring-measurement). Briefly, the rings of both Asl-mCherry and various GFP tagged CP110/Cep97 markers were fitted with an elliptical annular Gaussian profile, obtaining fit parameters for the major and minor axis. Centrioles that had Asl and CP110/Cep97 ring eccentricity (major:minor axis ratio) of less than 1.2 were considered well oriented. The width of the resulting Gaussian fit for CP110/Cep97 was then measured.

### Immunoblotting

Immunoblotting was carried out as described previously (Aydogan et al., 2018). For all blots, 15 staged early embryos were boiled in sample buffer and loaded in each lane. Primary antibodies used in this study are as follows: rabbit anti-CP110 (Franz et al., 2013), rabbit anti-Cep97 (Dobbelaere et al., 2020), rabbit anti-Cnn (Dobbelaere et al., 2008) and mouse anti-Actin (Sigma); all primary antibodies were used at 1:500 dilution in blocking solution (Aydogan et al., 2018). Primary antibody incubation period was 1h. Secondary antibodies used in this study are as follows: HRPO-linked anti-mouse or anti-rabbit IgG (both GE Healthcare); both the secondary antibodies were used at 1:3000 dilution in blocking solution (Aydogan et al., 2018). To estimate the relative expression of transgenes we performed serial dilution blots to compare with the levels of endogenous proteins.

The quantification of Western blots was carried out in ImageJ. Briefly, the endogenous CP110 or Cep97 signal in all conditions was selected by using the *Rectangle* tool to create a region of interest (ROI). The detected signals in each lane were plotted using *Plot Lanes* option from the *Gels* tab. The area under the curve for each lane was then calculated and exported to GraphPad Prism 8. The same size ROI was used to repeat the process for the loading control (the endogenous Actin signal) in each lane. The CP110 or Cep97 signals were normalized to the signal of the loading control in each lane, then to the respective mean value of CP110 or Cep97 signals overall.

### Fluorescence Correlation Spectroscopy (FCS)

FCS measurements were obtained as previously described (Aydogan et al., 2020). Every measurement consisted of 6x 10 sec point recordings, which were acquired around the centriolar plane (near the embryo cortex) at the beginning of nuclear cycle 12. The laser power was kept constant at 6.31 μW. All recordings were fitted with 8 previously described diffusion models (Aydogan et al., 2020) within the boundaries of 4x10^-4^ – 2.1x10^3^ ms using *FoCuS-point* (Waithe et al., 2016). The diffusion parameters were restricted to a minimal mean residence time of 0.7 ms, and the anomalous subdiffusion α and the spatial description of the excitation volume *AR* were kept constant at 0.7 and 5, respectively. The preferred model was chosen based on the Bayesian Information Criterion (BIC), which in our case was one diffusing species with one blinking and one triplet state of the fluorophore (Model 4). Afterwards, the cytoplasmic concentrations were corrected for the background noise (through point FCS measurements in n=21 WT embryos), and a ROUT outlier test (Q = 1%) was performed on all 10-sec long concentration measurements. Only the measurements with four or more recordings were kept for statistical analysis.

### Peak Counting Spectroscopy (PeCoS)

PeCoS measurements were obtained as previously described (Aydogan et al., 2020). 180 sec long recordings were made at the same position and the same nuclear cycle stage as the FCS measurements. In addition to the measurements of embryos expressing Plk4-GFP under its own endogenous promoter, 12 control measurements were obtained from embryos expressing Asl-mKate2, which were used to determine the background auto- fluorescence. The subtraction of their background (=“Mean+7*SD”) resulted in an average peak count of 4.5 (which fulfilled the requirement of less than 5 peaks per 180 sec-long control recording), and it was therefore chosen as background threshold for all *in vivo* measurements. The ROUT outlier test (Q = 1%) was performed before further statistical tests were applied.

### Quantification and statistical analysis

All the details for quantification, statistical tests, n numbers, definitions of center, and dispersion and precision measures were either described in the main text, relevant figure legends or in *Materials and Methods* section.

Significance in statistical tests was defined by *p*<0.05. To determine whether the data values came from a Gaussian distribution, D’Agostino-Pearson omnibus normality test was applied. GraphPad Prism 7 or 8 were used for all the modeling and statistical analyses, unless otherwise stated.

## Supplementary Tables

**Table S1:** D. melanogaster alleles used in this study.

| Allele* | Source (reference #) | ID |
| --- | --- | --- |
| Ubq-GFP-Cep97 | (Dobbelaere et al., 2008) | FlyBase ID: FBal0343980 |
| Ubq-CP110-GFP | (Dobbelaere et al., 2008) | FlyBase ID: FBtp0092320 |
| Ubq-CP110 | This paper | N/A |
| Ubq-Cep97 | This paper | N/A |
| Cep97-GFP | (Dobbelaere et al., 2020) | FlyBase ID: FBal0362412 |
| CP110-GFP | This paper | N/A |
| Plk4-GFP | (Aydogan et al., 2018) | FlyBase ID: FBal0343977 |
| Ubq-RFP-Cep97 | (Dobbelaere et al., 2008) | N/A |
| Plk4<sup>Aa74</sup> | (Aydogan et al., 2018) | FlyBase ID: FBab0049012 |
| CP110Δ | (Franz et al., 2013) | FlyBase ID: FBal0294119 |
| Cep97Δ | (Dobbelaere et al., 2020) | FlyBase ID: FBal0362411 |
| Asl-mCherry | (Conduit et al., 2015) | FlyBase ID: FBal0343645 |
| Ubq-Cep97-GFP | (Dobbelaere et al., 2008) | N/A |
| Sas-6-GFP | (Aydogan et al., 2018) | FlyBase ID: FBtp0131375 |
| Plk4 | (Aydogan et al., 2018) | FlyBase ID: FBal0343978 |
| Plk4<sup>RKA</sup> | (Aydogan et al., 2018) | FlyBase ID: FBtp0131379 |
| Asl-mKate2 | (Aydogan et al., 2020) | FlyBase ID: FBal0366991 |
| asl<sup>B46</sup> | (Baumbach et al., 2015) | FlyBase ID: FBal0343439 |
| CycB<sup>2</sup> | (Jacobs et al., 1998) | FlyBase ID: FBal0094855 |

\*The alleles listed here were expressed under their endogenous promoters unless otherwise specified.

**Table S2:** D. melanogaster strains generated and/or used in this study.

| Strain Genotype | Tissue | Type of experiment |
| --- | --- | --- |
| Ubq-GFP-Cep97 / + | Embryo | Confocal microscopy; Western blot; Fluorescence correlation spectroscopy |
| Ubq-CP110-GFP / + | Embryo | Confocal microscopy; Western blot; Fluorescence correlation spectroscopy |
| Cep97-GFP / + | Embryo | Confocal microscopy; Western blot |
| CP110-GFP / + | Embryo | Confocal microscopy; Western blot |
| Ubq-CP110-GFP / +; Plk4<sup>Aa74</sup> / + | Embryo | Fluorescence correlation spectroscopy |
| Ubq-GFP-Cep97/ +; +/+; Plk4<sup>Aa74</sup> / + | Embryo | Fluorescence correlation spectroscopy |
| Plk4-GFP / + | Embryo | Confocal microscopy; Peak counting spectroscopy |
| CP110Δ / CP110Δ; Plk4-GFP / + | Embryo | Confocal microscopy; Peak counting spectroscopy |
| Plk4-GFP / Ubq-CP110 | Embryo | Confocal microscopy; Peak counting spectroscopy |
| Plk4-GFP, Cep97Δ / Cep97Δ | Embryo | Confocal microscopy; Peak counting spectroscopy |
| Plk4-GFP / Ubq-Cep97 | Embryo | Confocal microscopy; Peak counting spectroscopy |
| Asl-mCherry / Ubq-CP110-GFP | Embryo; wing discs | 3D-SIM; Airy-scan super resolution microscopy |
| Ubq-GFP-Cep97 / +; Asl-mCherry / + | Embryo; wing discs | 3D-SIM; Airy-scan super resolution microscopy |
| Asl-mCherry / +; Ubq-Cep97-GFP / + | Wing discs | 3D-SIM |

**Table S2 Cont'd: *D. melanogaster* strains generated and/or used in this study.**
| Strain Genotype | Tissue | Type of experiment |
| --- | --- | --- |
| *Oregon-R* (Wild-type strain) | Embryo | Western Blot; Hatching assay; Fluorescence correlation spectroscopy |
| CP110Δ / CP110Δ | Embryo | Western blot; Hatching assay |
| Ubq-CP110 / + | Embryo | Western blot; Hatching assay |
| Cep97Δ / Cep97Δ | Embryo | Western blot; Hatching assay |
| Ubq-Cep97 / + | Embryo | Western blot; Hatching assay |
| Ubq-CP110-GFP / Ubq-RFP-Cep97 | Embryo | Confocal microscopy |
| Sas-6-GFP / + | Embryo | Confocal microscopy; Airy-scan super resolution microscopy |
| CP110Δ / CP110Δ; Sas-6-GFP / + | Embryo | Confocal microscopy; Airy-scan super resolution microscopy |
| Sas-6-GFP / Ubq-CP110 | Embryo | Confocal microscopy |
| Sas-6-GFP, Cep97Δ / Cep97Δ | Embryo | Confocal microscopy; Airy-scan super resolution microscopy |
| Sas-6-GFP / Ubq-Cep97 | Embryo | Confocal microscopy |
| Asl-mCherry / Sas-6-GFP | Embryo | Airy-scan super resolution microscopy |
| CP110Δ / CP110Δ; Asl-mCherry / Sas-6-GFP | Embryo | Airy-scan super resolution microscopy |
| Asl-mCherry, Cep97Δ / Sas-6-GFP, Cep97Δ | Embryo | Airy-scan super resolution microscopy |
| Ubq-GFP-Cep97 / +; Plk4 / + | Embryo | Fluorescence correlation spectroscopy |

**Table S2 Cont'd: *D. melanogaster* strains generated and/or used in this study.**
| Strain Genotype | Tissue | Type of experiment |
| --- | --- | --- |
| Ubq-GFP-Cep97 / +; + /+;<br>Plk4<sup>RKA</sup> / + | Embryo | Fluorescence correlation spectroscopy |
| Ubq-CP110-GFP / Plk4 | Embryo | Fluorescence correlation spectroscopy |
| Ubq-CP110-GFP / +; Plk4<sup>RKA</sup> / + | Embryo | Fluorescence correlation spectroscopy |
| Plk4<sup>Aa74</sup> / + | Embryo | Western blot |
| Plk4 / + | Embryo | Western Blot |
| Plk4<sup>RKA</sup> / + | Embryo | Western blot |
| Asl-mKate2, asl<sup>B46</sup> / + | Embryo | Peak counting spectroscopy |
| Plk4-GFP / Ubq-RFP-Cep97 | Embryo | Confocal microscopy |
| Ubq-CP110-GFP / CycB<sup>2</sup> | Embryo | Confocal microscopy |
| Ubq-GFP-Cep97 / +; CycB<sup>2</sup> / + | Embryo | Confocal microscopy |

**Table S3:**
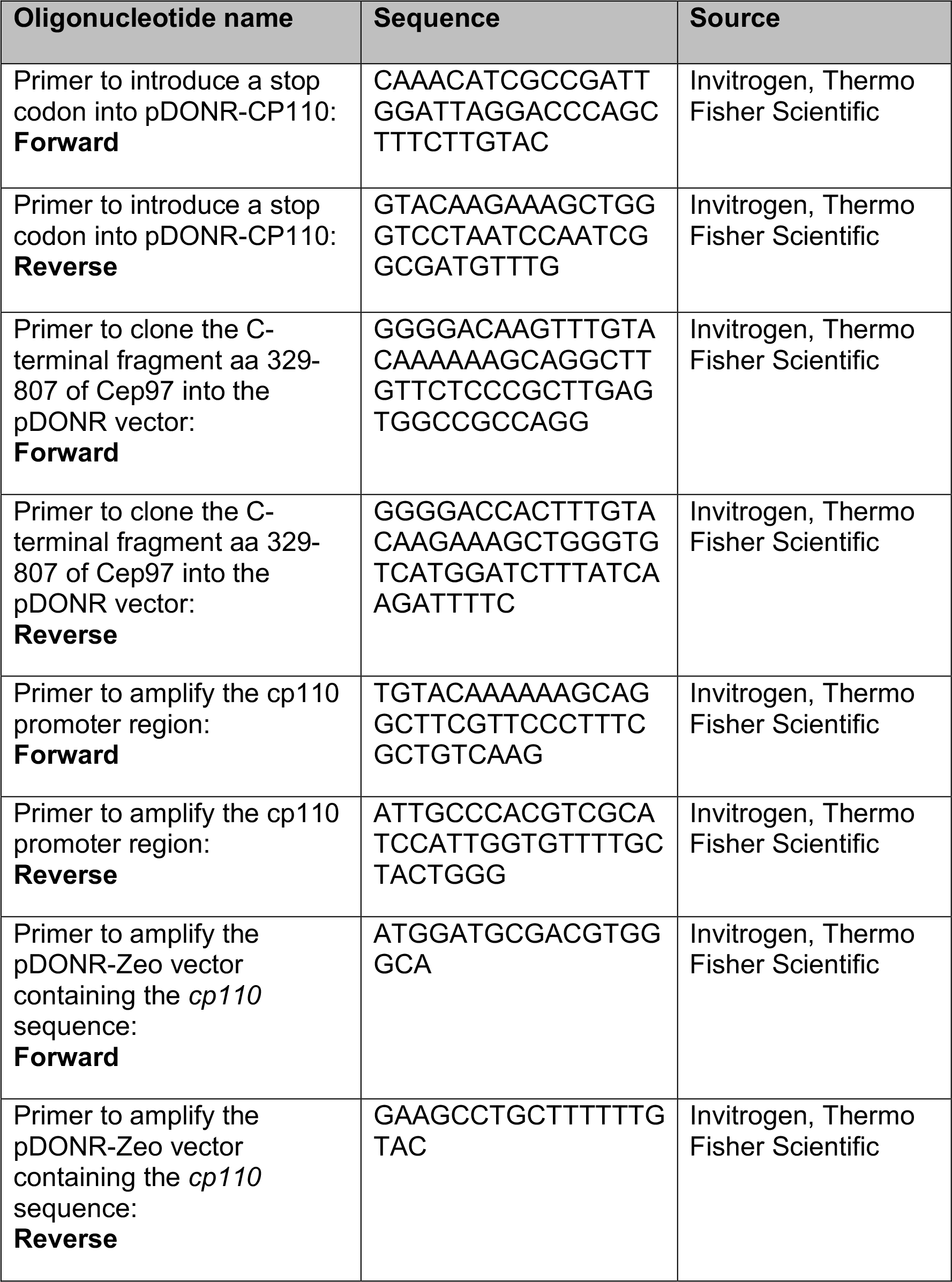
Oligonucleotides used in this study.

## Supplementary figure legends

**Figure S1.**
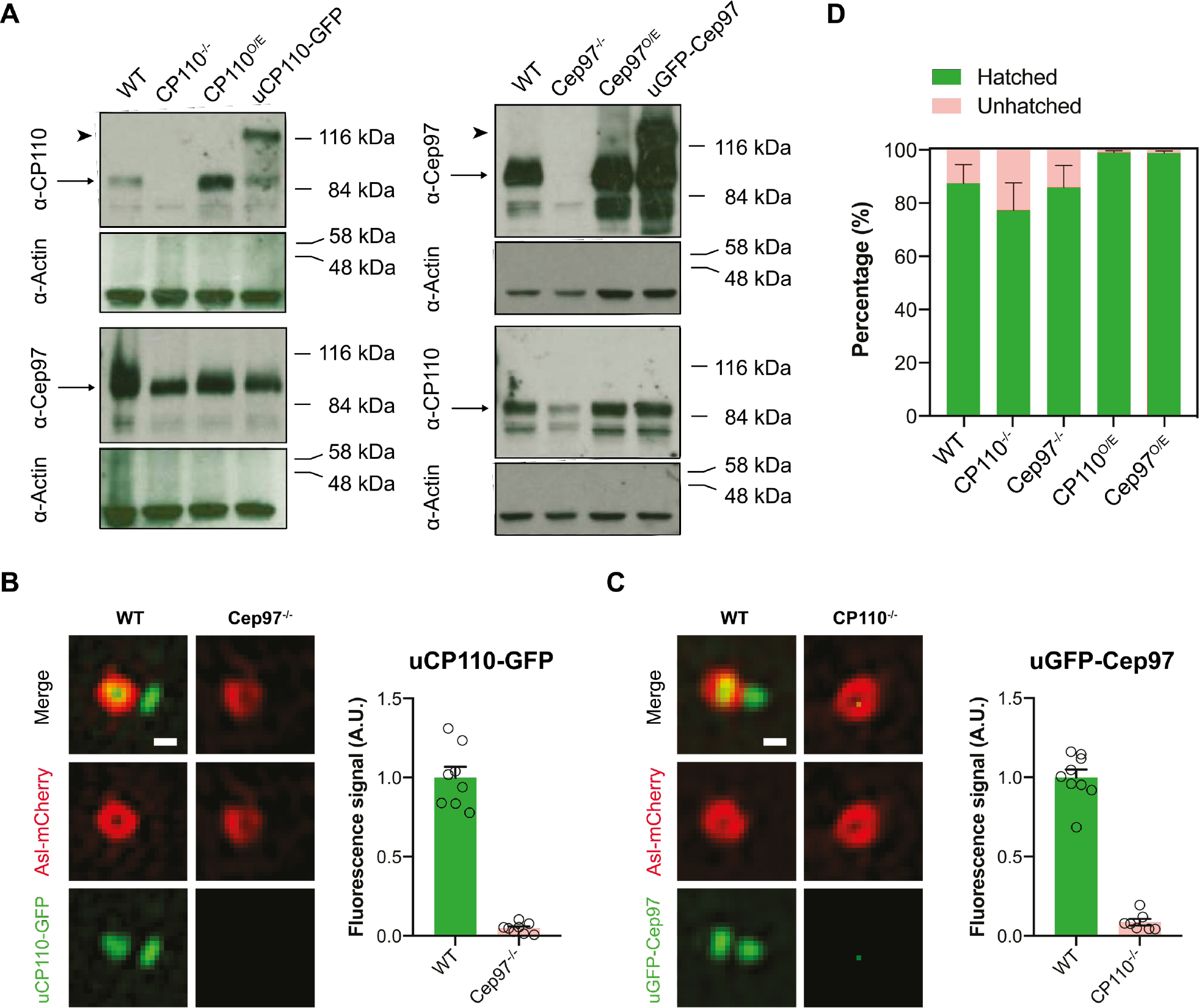
CP110 and Cep97 are largely co-dependent for their centriolar localisation and partially co-dependent for their cytoplasmic stability. **(A)** Western blots show the protein levels of CP110 and Cep97 in WT embryos, *CP110* or *Cep97* null mutant embryos (-/-), embryos overexpressing untagged CP110 or Cep97 from the *Ubq* promoter (O/E), or embryos overexpressing CP110-GFP or GFP-Cep97 from the Ubq promoter (u). Actin is shown as a loading control. Representative blots are shown from three technical repeats. Note that there appears to be slightly less Cep97, which is less smeared, in the absence of CP110, while there is clearly less CP110 in the absence of Cep97. **(B and C)** Airy-scan micrographs shows the centriole localisation of either uCP110-GFP (*green*, B) in WT and *Cep97^-/-^* embryos or uGFP-Cep97 (*green*, C) in WT and *CP110^-/-^* embryos; Asl-mCherry (*red*) labels the mother centrioles (Scale bar=0.2 μm). Bar charts quantify the centriolar levels (Mean±SD) of uCP110-GFP or uGFP-Cep97 in these embryos. For this quantification, the ten centriole pairs with the brightest Asl-mCherry were selected in each embryo (as CP110 or Cep97 levels could not be used to reliably identify the centrioles in the mutant embryos) and the centriolar levels of uCP110-GFP or uGFP-Cep97 was measured at 20 minutes into interphase of cycle 14. We analysed cycle 14 embryos in this experiment because CP110 and Cep97 centriolar levels rise to a steady plateau in the first 5-10mins of the extended interphase period in this cycle (rather than dropping as the embryos prepare to enter mitosis as in the earlier cycles), so centriolar fluorescence is normally at a constantly high level at this stage. N≥7 embryos, n=10 centrioles per embryo. **(D)** Bar chart indicates the embryo hatching frequency in wild type flies (*Oregon R*) or in flies of the indicated genotypes.

**Figure S2.**
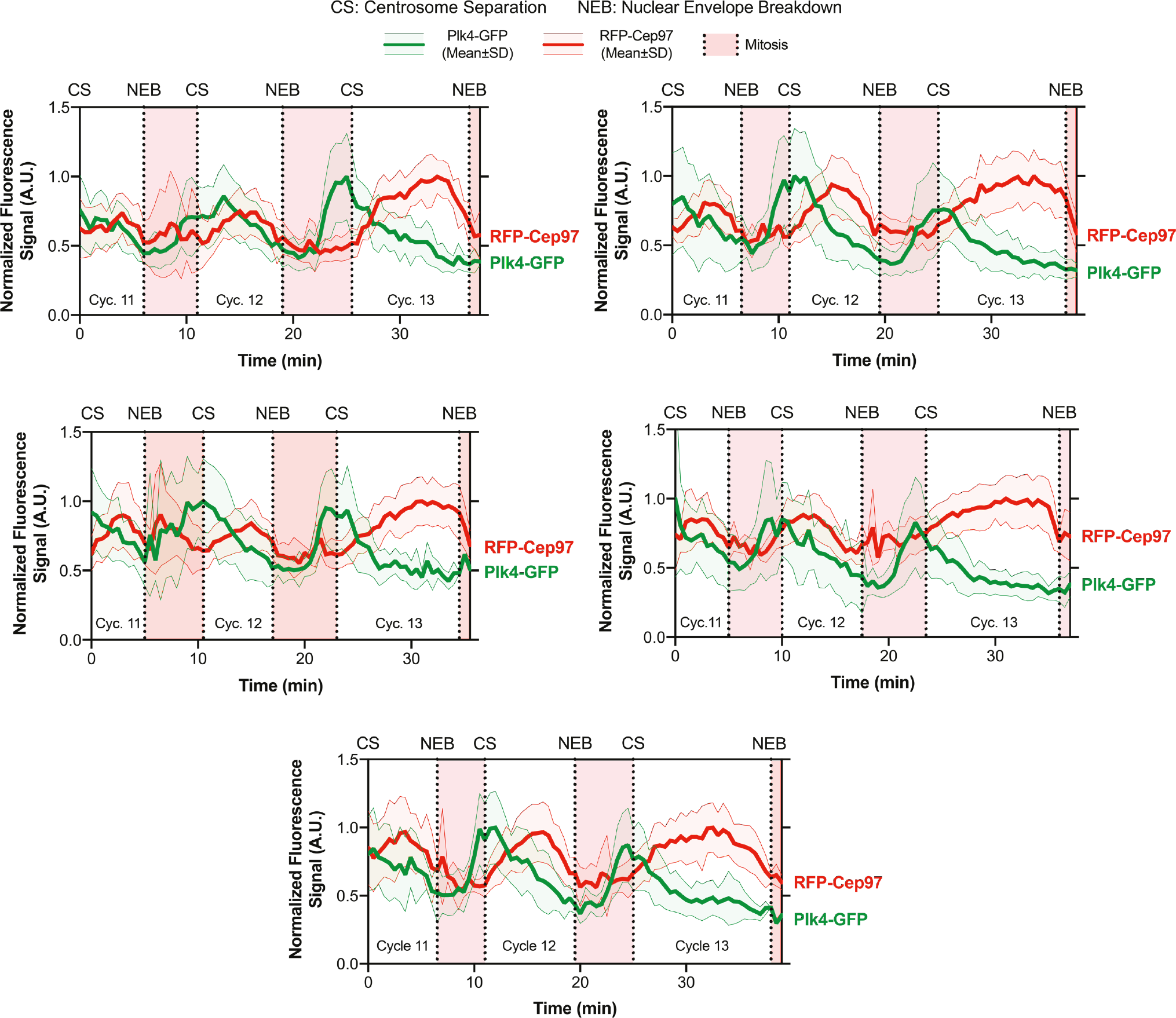
The centriolar recruitment of Cep97 is largely out of phase with the centriolar recruitment of Plk4. Graphs quantify the centriolar fluorescence levels (Mean±SD) of Plk4-GFP (*green*) and uRFP-Cep97 (*red*) co-expressed in five individual embryos analysed during nuclear cycles 11-13. CS= Centrosome Separation, NEB=Nuclear Envelope Breakdown. An average of n=41 centrioles were tracked per embryo.

**Figure S3.**
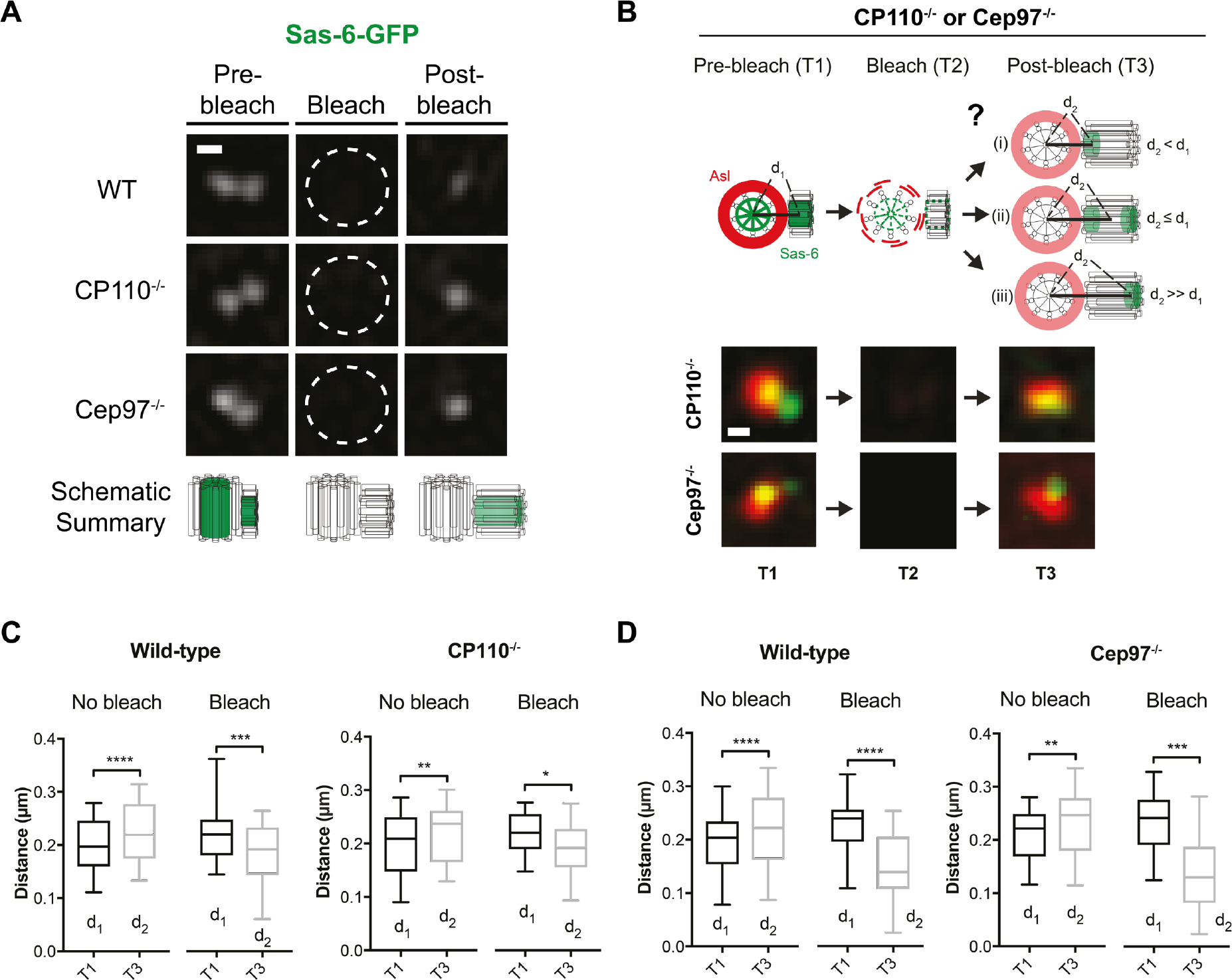
The centriole cartwheel continues to grow preferentially from the proximal-end of daughter centrioles in CP110^-/-^ and Cep97^-/-^ embryos. **(A)** Micrographs show a 3D-SIM-FRAP analysis of Sas-6-GFP dynamics in WT, *CP110^-/^*^-^ and *Cep97^-/-^* embryos. For each condition, a 3D-SIM image of a centriole pair was acquired in early/mid S-phase (Pre-bleach). The centrioles were subsequently photobleached (Bleach), and a 3D-SIM image was acquired 1 min after photobleaching (Post-bleach). These observations demonstrate that Sas-6-GFP continues to be incorporated exclusively into the growing daughter centriole even in the absence of CP110 or Cep97. Scale bar=0.2 μm. N≥8 embryos per group, n=3 centriole pairs on average per embryo. Schematics below each micrograph illustrate our interpretation of the FRAP experiments. **(B)** Schematic illustrates the photobleaching assay previously used to show that Sas-6-GFP preferentially incorporates into the proximal-end of growing daughter cartwheels (outcome [i]) (Aydogan et al., 2018). We used the same assay to test whether this was also the case in *CP110^-/-^* and *Cep97^-/-^* embryos. (Lower panel) Airy-scan super resolution micrographs show representative centriole images during pre-bleach (T1), bleach (T2) and post-bleach (T3) stages of the FRAP experiment in CP110^-/-^ and Cep97^-/-^ embryos simultaneously expressing Asl-mCherry and Sas-6- GFP. Scale bar=0.2 μm. **(C and D)** Box and whisker plots show the pre- and post-bleach distance (d1 and d2, respectively) between Asl-mCherry on the mother centriole and the newly incorporating Sas-6-GFP on the growing daughter centriole in *CP110^-/-^* (C) or *Cep97^-/-^* (D) embryos compared to WT controls. In the *No bleach*control experiment, d2 > d1 for all conditions, reflecting the growth of the daughter centriole between T1 and T3. In the *Bleach* experiment, d2 << d1 for all conditions, indicating that Sas-6-GFP continues to incorporate only into the proximal-end of the centrioles in the absence of CP110 or Cep97. N≥11 embryos per condition; n≥16 centriole pairs for *No Bleach* and *Bleach* groups each. Midlines represent the median, whiskers (error bars) mark the minimum to maximum, and bottom/top of the boxes indicate the first/third quartile of the distribution, respectively. Statistical significance was assessed using a paired *t* test (*, P<0.05; **, P<0.01; ***, P<0.001; ****, P<0.0001).

## Supplementary videos and captions

Movie S1. Monitoring the centriolar dynamics of uCP110-GFP in a

***Drosophila* embryo.**

Time-lapse video of an embryo expressing uCP110-GFP, observed on a spinning-disk confocal microscope through nuclear cycles 11-13. The movie is a maximum-intensity projection that has been photo-bleach corrected, but not background subtracted for visual clarity. Time (min:sec) is shown at the top left, and the developmental stage of the embryo is indicated at the bottom left.

Movie S2. Monitoring the centriolar dynamics of uGFP-Cep97 in a

***Drosophila* embryo.**

Time-lapse video of an embryo expressing uGFP-Cep97, observed on a spinning-disk confocal microscope through nuclear cycles 11-13. The movie is a maximum-intensity projection that has been photo-bleach corrected, but not background subtracted for visual clarity. Time (min:sec) is shown at the top left, and the developmental stage of the embryo is indicated at the bottom left.

Movie S3. Monitoring the centriolar dynamics of uCP110-GFP and uRFP- Cep97 simultaneously in the same embryo.

Time-lapse movie of an embryo expressing uCP110-GFP and uRFP-Cep97, observed on a spinning-disk confocal microscope through nuclear cycles 11- 13. The movie is a maximum-intensity projection that has been photo-bleach corrected, but not background subtracted for visual clarity. Time (Min:Sec) is shown at the top left, and the developmental stage of the embryo is indicated at the bottom left.

